# Understanding the Human Brain using Brain Organoids and a Structure-Function Theory

**DOI:** 10.1101/2020.07.28.225631

**Authors:** Gabriel A. Silva, Alysson R. Muotri, Christopher White

## Abstract

A basic neurobiology-clinical trial paradigm motivates our use of constrained mathematical models and analysis of personalized human-derived brain organoids toward predicting clinical outcomes and safely developing new therapeutics. Physical constraints imposed on the brain can guide the analyses an interpretation of experimental data and the construction of mathematical models that attempt to make sense of how the brain works and how cognitive functions emerge. Development of these mathematical models for human-derived brain organoids offer an opportunity for testing new hypotheses about the human brain. When it comes to testing ideas about the brain that require a careful balance between experimental accessibility, manipulation, and complexity, in order to connect neurobiological details with higher level cognitive properties and clinical considerations, we argue that fundamental structure-function constraints applied to models of brain organoids offer a path forward. Moreover, we show these constraints appear in canonical and novel math models of neural activity and learning, and we make the case that constraint-based modeling and use of representations can bridge to machine learning for powerful mutual benefit.

## 1 Introduction

Everything the human brain is capable of is the product of a complex symphony of interactions between many distinct signaling events and myriad of individual computations. The brain’s ability to learn, connect abstract concepts, adapt, and imagine, are all the result of this vast combinatorial computational space. Even elusive properties such as self-awareness and consciousness presumably owe themselves to it. This computational space is the result of thousands of years of evolution, manifested by the physical and chemical substrate that makes up the brain - its ’wetware’.

Physically, the brain is a geometric spatial network consisting of about 86 billion neurons and 85 billion non-neuronal cells [3, 12]. There is tremendous heterogeneity within these numbers, with many classes and sub-classes of both neurons and non-neuronal cells exhibiting different physiological dynamics and morphological (structural) heterogeneity that contribute to the dynamics. A seemingly uncountable number of signaling events occurring at different spatial and temporal scales interacting and integrating with each other is the source behind its complexity and the functions that emerge.

To get a sense of the sheer size of the numbers that make up this computational space, a first order approximation of the total number of synapses is roughly about ten quadrillion [8]. Individual neurons have been estimated to have order of magnitude 10,000’s to 100,000’s synapses per neuron. The exact number of synapses varies significantly during the course of a person’s lifetime and brain region being considered. For example, pyramidal neurons in the cortex have been estimated to have about 32,000 synapses [17], interneruons, which are the computational workhorses of the brain, between 2,200 and 16,000 [14], neurons in the rat visual cortex have about 11,000 per neuron [18], and neurons in other parts of the cortex in humans have roughly 7,200 synapses [28, 21].

Yet, despite a size that is intuitively difficult to grasp, this computational space is finite. It is limited by the the physical constraints imposed on the brain. The thesis we consider here is that taking advantage of these constraints can guide the analyses and interpretation of experimental data and the construction of mathematical models that aim to make sense about how the brain works and how cognitive functions emerge. Specifically, we propose that there exists a fundamental structure-function constraint that any theoretical or computational model that makes claims about relevance to neuroscience must be able to account for. Even if the intent and original construction of the model do not not take this constraint into consideration.

Testing candidate theoretical and computational models has to rely on the use of mathematics as a unifying language and framework [27]. This lets us build an engineering systems view of the brain that connects low-level observations to functional outcomes. It also lets us generalize beyond neuroscience. For example, understanding the algorithms of the brain separately from the wetware that impliment them will allow these algorithms to be integrated into new machine learning and artificial intelligence architectures. As such, mathematics needs to play a central role in neuroscience, not a supporting one. To us this intersection between mathematics, engineering, and experiments reflects the future of neuroscience as a field.

There must also be plausible experimentally testable predictions that result from candidate models. A particular sensitivity is that the predictions of mathematical models may exceed the expressive capabilities of existing experimental models. This motivates the push for continuously advancing experimental models and the technical methods required to interrogate them. While the list to choose from across many different species and scales is long, and no experimental preparation of course is ever perfect, we argue that human-derived brain organoids offer what could be a unique model of the human brain for this purpose. In particular when it comes to testing ideas about the brain that require a careful balance between experimental accessibility, manipulation, and complexity in order to bridge neurobiological details with higher level cognitive properties in the intact human. This includes clinical considerations of relevance to health and disease. Or the application of mathematical models and algorithms about the brain to machine learning and artificial intelligence. Abstraction (for generalization) and validation, through human organoids to humans, and models to machines, both support a systems-approach building on constraints while modeling behavior.

The rest of this paper we will explore these ideas in detail. We will start by exploring how brain organoids from individual research subjects and patients offer an individualized opportunity for personalized neuroscience that can occur in parallel with cognitive studies or clinical trials. We then discuss the fundamental structure-function constraint and look at theoretical models that explicitly or implicitly take this constraint into account. Finally, we also review experimental results where the constraint anchors the interpretation of data.

## 2 Human-derived brain organoids as an experimental model

First developed by Yoshi Sassai and colleagues [15, 19, 20, 24, 25], neural structures can be derived from human pluripotent stem cells. Brain organoids are pin-sized three dimensional self-organizing structures composed of roughly 2.5 million neural cells. They can be generated from human pluripotent stem cells, either embryonic stem cells directly or derived from human blastocysts, and from induced pluripotent stem cells reprogrammed from somatic donor cell types. Beginning with terminally differentiated cells, such as skin fibroblasts, genetic and cellular reprogramming is carried out by carefully orchestrated exposure to specific sets of transcription factors and chemical environments [2, 29, 30]. The outcome is the de-differentiation of the initial cells back to an embryonic-like pluriopotent stem cell state, with subsequent guided differentiation into different types of neurons and other neural cells. The recapitulation of the developmental genetic program -or at least certain aspects of it -produces three dimensional structures that spontaneously self-organize at a cellular scale into anatomical structures that resemble aspects of the prenatal brain. For example, there is a distinctly defined cortical plate on its surface, with a central cavitation resembling a ventricle. The molecular composition of the constituent cells include markers for mature neurons and glia, although markers for progenitors and immature cells are also present. It was recently shown that if the culture conditions are appropriate, sufficient growth and maturation of organoids results in them being able to display local field potential (LFP) like electrophysiological activity. Organoid LFP’s are sufficiently similar in their firing frequency and patterns to prenatal human brains that a machine learning algorithm trained on features of prenatal EEG’s can accurately identify relative age matched organoids when shown the organoid LFP’s [2].

### 2.1 Bridging basic neuroscience to clinical and cognitive research

Human-derived brain organoids reflect a personalized model of the brain unqique to each individual. Because of this, they could provide an opportunity to bridge biological experiments and computational models with behavioral, cognitive, and clinical studies in humans that no other experimental model is able to achieve. Organoids derived from neurotypical individuals or patients can be experimentally studied in parallel with cognitive experiments or clinical trials being carried out in the same individuals. They can potentially put molecular, cellular and physiological scale experiments distinct to each individual in context with cognitive and clinical studies that the subjects or patients are participating in. This would be invaluable for the investigation of physiologically complex genetically based neurodevelopmental and neurological disorders such as autism spectrum disorder and epilepsy (Figure 1). Presumably, during the course of the cellular reprogramming, the genetic program responsible for the development of that specific individual is reawakened and recapitulated. Absent epigenetic, environmental, and myriad other factors that actually shape the development of the brain - clearly an uncontrolled but unavoidable limitation of any such model -brain organoids may offer an opportunity to study and understand key aspects of the neurobiology initiated by each person’s unique genetic program. A basic science-clinical experimental design paradigm.

**Figure 1:**
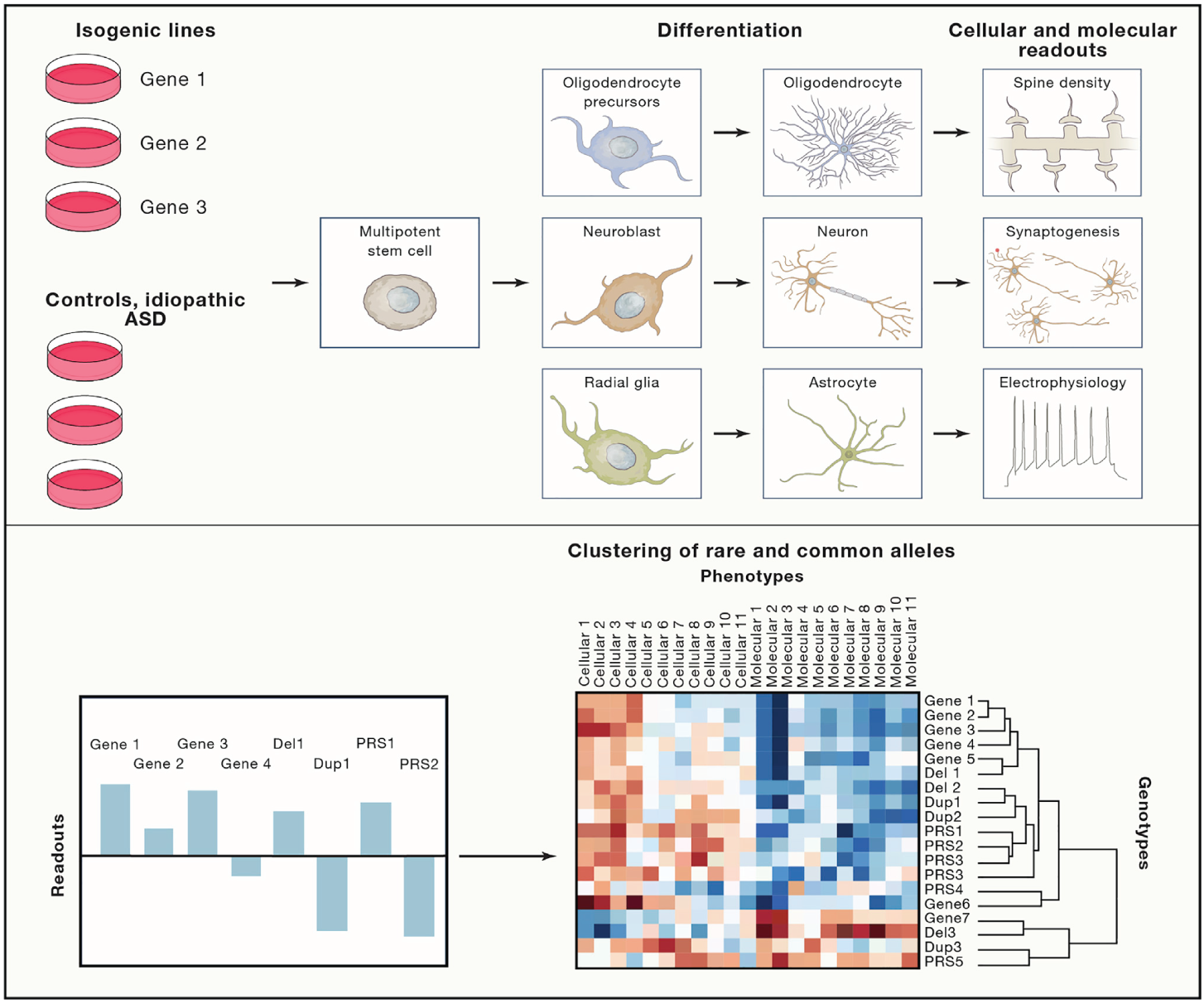
Identifying core gene sets that regulate neurodevelopment based on trait correlations. The characterization of genotype-trait correlations in brain organoid models can define sets of genes and copy number variations that share common phenotype profiles. These profiles could reflect conditions associated with common neuronal function. For example, the effects of multiple gene mutations and copy number variations can be tested in relation to isogenic controls across a series of cellular assays. Subsequent analysis allow genes, copy number variations, and common variants to be clustered into discrete groups. Comparing patterns of trait correlations in organoid models versus clinical phenotype data (not shown) could help to identify clinical subtypes that share common neuropathologies. Reproduced from [13].

In fact, we are currently pursuing this approach in a study at the University of California San Diego focused on the effects of cannabidiol (CBD) on autism [32], A key objective of this study is to test a hypothesis in patient-specific derived brain organoids that there is a breakdown in information (signaling) transfer efficiencies in neurons and neuronal networks in autistic individuals. We are investigating this by attempting to measure deviations in a calculation we call the refraction ratio. The refraction ratio is derived from theoretical considerations of the fundamental structure-function constraint we discuss below. We will then compute the refraction ratio again following bath exposure to different concentrations of CBD and compare it to the data prior to treatment. This is all occurring in parallel while the patients themselves are participating in a CBD clinical trial. One possibility we are testing is if there are aspects of a potential clinical response that can be predicted by the refraction ratio-organoid basic studies. It is conceivable that in future clinical studies parallel *in vitro* studies with brain organoids are used to optimize testing or dosing conditions, for example, or to provide physiological context to observed clinical effects. This dual basic neurobiology and computational neuroscience analysis in parallel with a clinical study or individualized cognitive experiments cannot realistically be done in any other experimental model. It requires a degree of personalization of the uniqueness of each individual’s brain, or at least aspects of it, that no animal model or ’generalized’ model of the brain can achieve. At the same time, brain organoids seem to be sufficiently complex that they provide a real test bed for systems neuroscience theoretical and computational models specific to the individual subject or patient.

### 2.2 Open questions

To be sure, there are many unanswered questions about exactly how appropriate organoids are as a model of the brain. Brain organoids are not brains and do display a number of limitations. For example, they are not vascularized, do not have all cell types present, and retain populations of immature neurons with confused identities. There are also a number of technical challenges that must be addressed in order to study them properly, such as how to image their connectome given their size, and how best to do three dimensional electrophysiology given their geometry. But answering these open questions and continuing to improve the experimental methods in order to assess the utility of organoids as a sufficiently complex model of the human brain is very much worth the pursuit, and at the forefront of this research. There is a high level of excitement at the prospect of them being a unique developmental and physiological model of the brain.

Scientifically, a number of questions about the impact of the structure-function constraint on neural development and functions specific to the human brain motivate much of our work. Yet, we can only realistically address these questions using the brain organoid model, or something like it. On the one hand, it is unclear how generalizable answers to our questions derived from experimental models from other species would be. On the other, the technical requirements associated with the measurements and manipulations necessary to answer these questions cannot be done in intact humans. For example, does the genetic program responsible for the development of the brain support a connectivity structure of the network that is stochastically but purposefully wired? In other words, are the geometric and topological properties of the brain purposely designed, or are they just random? A naive interpretation of what we mean by ‘purposely built’ is a structural network designed to sustain inherent and recurrent activity in response to a stimulus (either external, i.e sensory, or internally generated), by satisfying the dynamic constraints of non-zero and non-saturated activity. Presumably, it is this functional regime that supports sustained recurrent activity resulting in rich dynamical repertoires in order to achieve a meaningful neurophysiological operating range. How is this designed in the brain to support meaningful neural dynamics? Below we give an example of how easily the ’wrong’ geometry and connectivity in a network can kill dynamic activity, but do so in a simulation of a biological neural network [6]. We also discuss results that suggest that in the worm C. elegans at least, the wiring of the organism’s connectome seems to have evolved to intentionally support sustained recurrent activity [10]. But such a basic theoretical hypothesis about the operation of the brain, grounded in fundamental ideas about its physical structure and dynamics, cannot be directly tested in humans. Brain orgaoids offer the opportunity to bridge this gap.

## 3 A fundamental structure-function constraint

### 3.1 A fundamental constraint imposed by neural structure and dynamics

Any theoretical or computational model of the brain, regardless of scale, must be able to account for the constraints imposed by the physical structures that bound neural processes. This is a reasonable requirement if the claims being made by a model are that the brain actually works in the way its describing. The conditions that determine these physical constraints are informed by neurobiological data provided by experiments.

One very basic constraint is the result of the interplay between anatomical structure and signaling dynamics. The integration and transmission of information in the brain are dependent on the interactions between structure and dynamics across a number of scales of organization. This fundamental structure-function constraint is imposed by the geometry (connectivity and path lengths) of the structural networks that make up the brain at different scales, and the resultant latencies involved with the flow and transfer of information within and between functional scales. It is a constraint produced by the way the brain is wired, and how its constituent parts necessarily interact, e.g. neurons at one scale and brain regions at another.

The networks that make up the brain are physical constructions over which information must travel. The signals that encode this information are subject to processing times and signaling speeds (conduction velocities) that must travel finite distances to exert their effects - in other words, to transfer the information they are carrying to the next stage in the system. Nothing is infinitely fast. Furthermore, the latencies created by the interplay between structural geometries and signaling speeds are generally at a temporal scale similar to the functional processes being considered. So unlike turning on a light switch, where the effect to us seems instantaneous and we can ignore the timescales over which electricity travels, the effect of signaling latencies imposed by the biophysics and geometry of the brain’s networks cannot be ignored. They determine our understanding of how structure determines function in the brain, and how function modulates structure, for example, in learning or plasticity.

While a model or algorithm does not explicitly need to take this fundamental structure-function constraint into account, it must have an implicit inclusion of constraint dependencies in order to be relevant to neurobiology. In other words, additional ideas and equations should be able to bridge the model’s original mathematical construction to an explicit dependency on the structure-function constraint.

For example, a model that relies on the instantaneous coupling of spatially separable variables can never be a model of a process or algorithm used by the brain because there is simply no biological or physical mechanism that can account for such a required mathematical condition. The requirement that any realistic candidate model of a brain algorithm be able to accommodate the structure-function constraint is simply the result of the physical make up and biophysics of biological neural networks.

### 3.2 The fundamental structure-function constraint in three different models

To illustrate these ideas, we discuss the dependency of the fundamental structure-function constraint in three distinct examples. We chose these models in part because they capture different functional scales, from molecular and ionic considerations at the single cell level to cellular neural networks to a more abstract notion of life-long versus progressive learning. The first two models have an explicit dependency on the constraint, while the third is implicit and requires extending its mathematical construction to account for it. In this section we qualitatively summarize each example. The Appendix extends this discussion and includes explicit references to the relevant equations.

#### The Hodgkin Huxley model

Consider the canonical Hodgkin Huxley model of a neuron. The fundamental structure-function constraint is explicit in the model’s equations. Current flows in the cable equation are computed along the dominant one dimensional direction of the axon as a function of time. The membrane biophysics, due to the effects of the participating sodium and potassium ion channels, completely determines the passive spatial and temporal decay of potentials along the membrane. This is captured by the time and space constants, respectively. The space constant is the distance over which the potential has decayed a certain amount of its original amplitude. This is a critical consideration for the spatial and temporal summation of post synaptic potentials along the dendritic tree towards reaching the threshold potential necessary to trigger an action potential at the axon hillock. The membrane potential dynamically depends on the spatial variable and propagation latencies of ionic changes that determine signaling events.

In the solution to its equations, the variables that account for these changes ultimately determine the temporal kinetics of the action potential. If the membrane refractory state associated with the recovery of sodium channel inactivation is taken into consideration, the conduction velocity of the traveling action potential can be derived. This reflects the rate at which information can be communicated or transferred between neurons. We refer the reader to any of the standard presentations of the Hodgkin Huxley model for full details [1, 5, 9, 16].

#### Competitive-refractory dynamics model

As a second example, we (G.S.) recently described the construction and theoretical analysis of a framework (competitive-refractory dynamics model) derived from the canonical neurophysiological principles of spatial and temporal summation [26]. Like the Hodgkin Huxley model, the dependency of the model on the fundamental structure-function constraint is explicit in its construction and equations. In fact, the entire theory takes advantage of the constraints that the geometry of a network invokes on latencies of signal transfer on the network. But unlike Hodgkin Huxley it is not a biophysical model. It is a mathematical abstraction applicable to any physically constructible network.

The framework models the competing interactions of signals incident on a target downstream node (e.g. a neuron) along directed edges coming from other upstream nodes that connect into it. The model takes into account how temporal latencies produce offsets in the timing of the summation of incoming discrete events due to the geometry (physical structure) of the network, and how this results in the activation of the target node. It captures how the timing of different signals compete to ‘activate’ nodes they connect into. This could be a network of neurons or a network of brain regions, for example. At the core of the model is the notion of a refractory period or refractory state for each node. This reflects a period of internal processing or inability to react to other inputs at the individual node level. The model does not assume anything about the internal dynamics that produces this refractory state, which could include an internal processing time during which the node is making a decision about how to react. In a geometric network temporal latencies are due to the relationship between signaling speeds (conduction velocities) and the geometry of the edges on the network (i.e. edge path lengths).

A key result from considerations of the interplay between network geometry and dynamics is the refraction ratio -the ratio between the refractory period and effective latencies. This ratio reflects a necessary balance between how fast information (signals) are propagating throughout the network, relative to how long each individual node needs to internally process incoming information. It is an expression that describes a local rule (at the individual node scale) that governs the global dynamics of the network. We can then use sets of refraction ratios to compute the causal order in which signaling events trigger other signaling events in a network. This allows us to compute (predict) and analyze the overall dynamics of the network.

The last important consideration for our purposes here, is that a theoretical analysis of the model resulted in the derivation of a set of (upper and lower) bounds that constrained a mathematical definition of efficient signaling for a structural geometric network running the dynamic model. This produced a proof of what we called the optimized refraction ratio theorem, which turned out to be relevant to the analyses of experimental data, which we discuss in the next section.

#### Support from experimental results

The effects of the structure-function constraint can be seen in experimental results through analyses that make use of the competitive-refractory dynamics model. We have shown in numerical simulations that network dynamics can completely break down in geometric networks (such as biological neural networks) if there is a mismatch in the refraction ratio [6, 26]. In numerical simulation experiments, we stimulated a geometric network of 100 neurons for 500 ms with a depolarizing current and then observed the effects of modifying the refraction ratio (Figure 2). While leaving the connectivity and geometry (edge path lengths) unchanged, we tested three different signaling speeds (action potential conduction velocities) at 2, 20, and 200 pixels per ms. We then observed the network to see how it was able to sustain inherent recurrent signaling in the absence of additional external stimulation. At the lowest signaling speed, 2 pixels per ms, we saw recurrent low-frequency periodic activity that qualitatively resembled a central pattern generator. When we increased the speed to 20 pixels per ms, there was still some recurrent activity but it was more sporadic and irregular. At 200 pixels per ms however, there was no signaling past the stimulus period. All the activity died away. This is the direct consequence of a mismatch in the refraction ratio. When signals arrive to quickly, the neurons have not had enough time to recover from their refractory periods and the incoming signals do not have an opportunity to induce downstream activations. The activity in the entire network dies.

**Figure 2:**
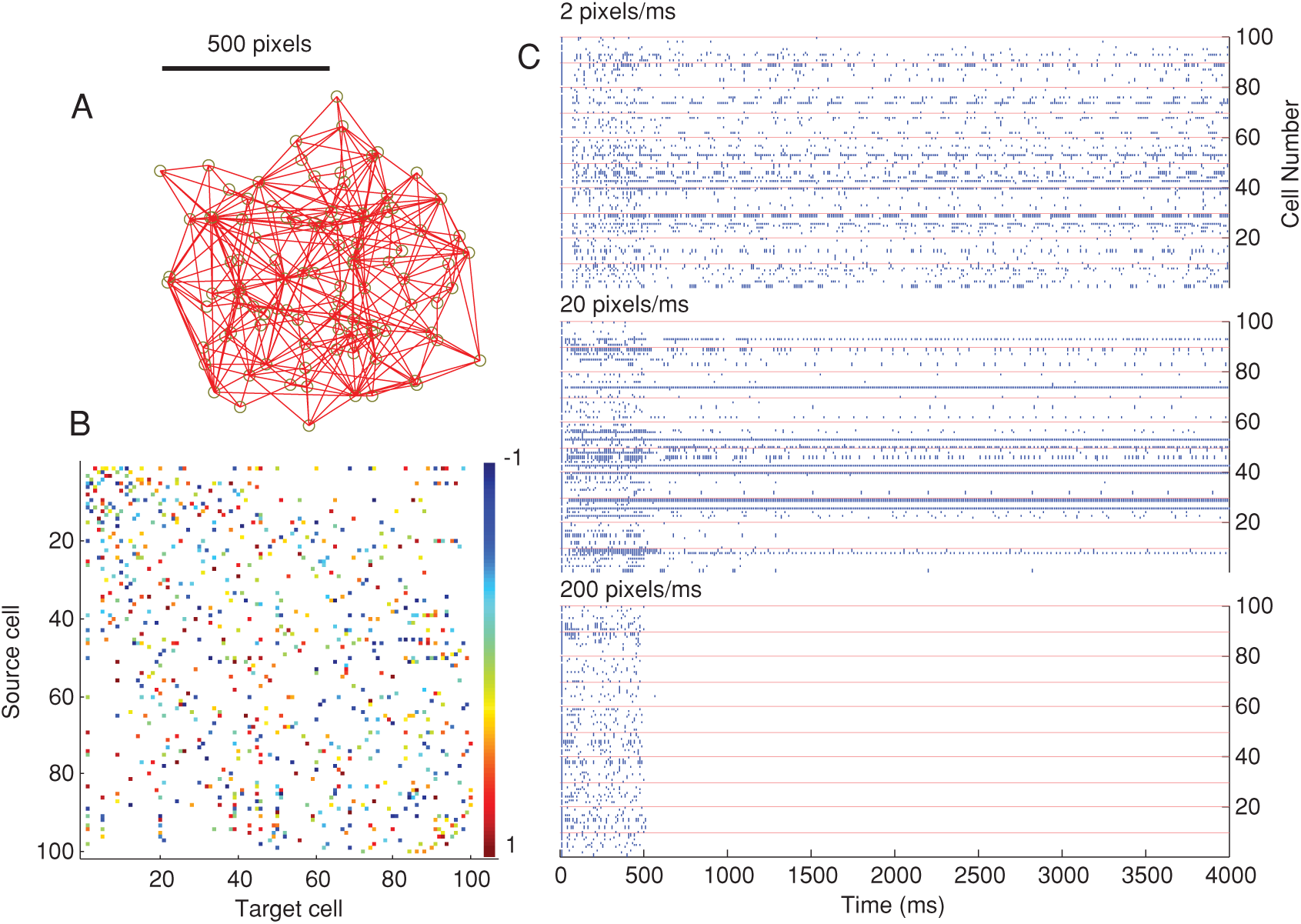
Effects of the structure-function constraint on the dynamics of a geometric network. **A**. We simulated a 100 node network of Izhikevitch neurons with a geometric construction (i.e. spatial coordinates of nodes) as shown in the figure. **B**. Functional connections were assigned random uniformly distributed weights between −1 and 1. **C**. Raster plots of the dynamics of the network for three different signaling speeds (conduction velocities). Reproduced from [6].

Using a refraction ratio analysis, we also showed that efficient signaling in the axon arbors of inhibitory Basket cell neurons depends on a trade-off between the time it takes action potentials to reach synaptic terminals (temporal cost) and the amount of cellular material associated with the wiring path length of the neuron’s morphology (material cost) [23]. Dynamic signaling on branching axons is important for rapid and efficient communication between neurons, so the cell’s ability to modulate its function to achieve efficient signaling is presumably important for proper brain function. Our results suggested that the convoluted paths taken by axons reflect a design compensation by the neuron to slow down action potential latencies in order to optimize the refraction ratio. The ratio is effectively a design principle that gives rise to the geometric shape (morphology) of the cells. Basket cells seem to have optimized their morphology to achieve an almost theoretical ideal of the refraction ratio (Figure 3).

**Figure 3:**
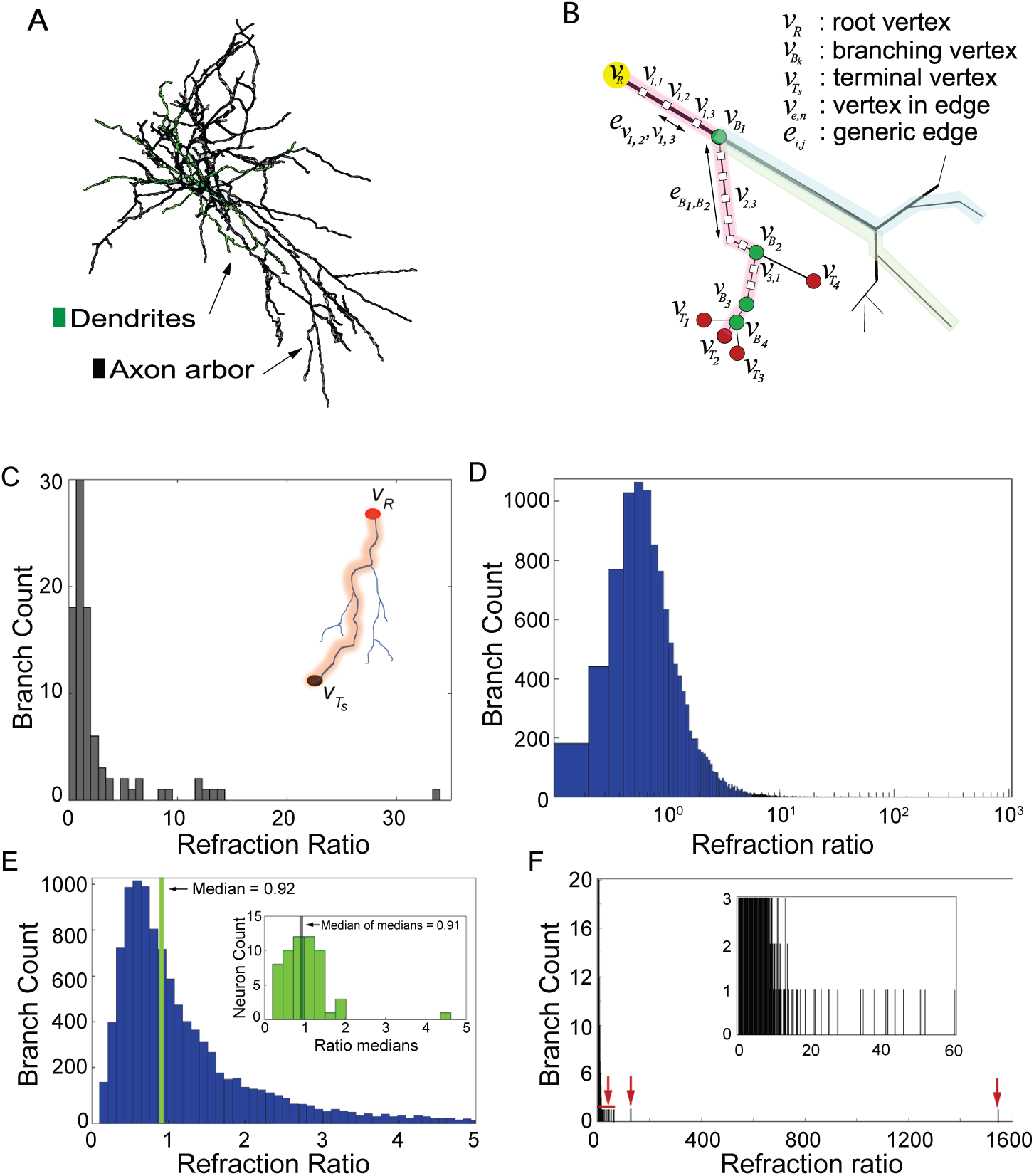
Refraction ratio analysis suggests Basket cell neurons optimize their morphology to the structure-function constraint. **A**. Three dimensional morphological reconstruction of one of the rat neocortical Basket cells from our data set. **B**. Model of the geometric network construction of an axon and its arborizations. We computationally approximated the edge path length integrals connecting the axon initial segment to a synaptic terminal **C**. Distribution of the refraction ratio for all branches forming the axon arbor for a single neuron. The inset shows an example of one axonal branch for this cell. The refraction ratio was computed independently for each axonal branch for all branches of the axon tree. The x-axes represents values of the computed refraction ratio, while the y-axes shows the number or count of individual axonal branches. **D**. Distribution of the refraction ratio for the full data set of 57 basket cells. Note the logarithmic scale in order to account for outlier ratio values. All axonal branches (11,575) from all the neurons were computed as statistically independent variables. **E**. The peak of the distribution shown in panel D occurred at 0.56, while the median had a value of 0.92. The median value of all medians (inset) computed for each of the 57 neurons had a value of 0.91 This allowed us to normalize for neurons that may have been contributing a disproportionate number of branches, thereby potentially skewing the median ratio for the entire population. The mean of the median ratio distribution had a value of 1.01 with 49 neurons (86 % of the entire data set) within one standard deviation. The theoretically derived ideal for efficient signaling in a geometric network is unity (see the refraction ratio theorem in [26]). (D) Distribution of the refraction ratio for the entire data set (same data shown in panel B) but with x-axis extending out to the full range of values in order to show the few outliers (indicated by the red arrows). Inset: zoomed in view of the main plot out to a refraction ratio of 60. Reproduced from [23].

We have also analyzed the signaling dynamics of the C. elegans connectome to infer a purposeful connectivity design to how the nervous system of the worm has evolved to be wired [10]. We simulated the dynamics of the C. elegans connectome using the competitive-refractory dynamics model that took advantage of the known spatial (geometric) connectivity of the worm. We studied the dynamics which resulted from stimulating a chemosensory neuron (ASEL) in a known feeding circuit, both in isolation and embedded in the full connectome. We were able to show that contralateral motor neuron activations in ventral and dorsal classes of motor neurons emerged from the simulations in such a way that they were qualitatively similar to rhythmic motor neuron firing pattern associated with the locomotion. Part of our interpretation of the results was that there is an inherent -and we proposed -purposeful structural wiring to the C. elegans connectome that has evolved to serve specific behavioral functions.

#### Progressive transfer learning model

The last example we will discuss is different than the first two models in that there is no explicit dependency on the fundamental structure-function constraint. But it is extendable to accommodate it. The model is a generalized approach for transfer learning developed by one of authors (C.W.) and colleagues [31], which we call progressive learning. This is an approach that supports transfer learning across tasks. In other words, given data from a new task, the goal is to use all the data from previous tasks to better learn new tasks then if we considered information from just the new task alone. This is a stronger requirement than simply avoiding what is called catastrophic forgetting, which implies that the progressive learning of additional new tasks does not degrade what you have learned previously. We defined a forward transfer efficiency and a backward transfer efficiency. Forward transfer efficiency measured a learning error given only information for the current task at hand. While backward transfer efficiency measured the learning error given access to all the prior learned information in addition to that for the current task.

The core of our argument relies on being able to evaluate a hypothesis that an observed input results in a given output. For the single task case, this hypothesis is decomposed into three parts, each with a successive purpose. Given an input, a transformer maps the input into an internal representation. Following the construction of an internal representation to the input, a voter maps the transformed input data into a posterior distribution on a response space. In the last step, a decider takes the information from the voters and maps the (now transformed input) to an output.

The model’s core insight and innovation is the construction of ensemble representations, or groupings, of learned transformations across tasks, which can then be used by deciders on new tasks. In other words, the transfer of learned information between different tasks contributes to learning new tasks. The decider combines the votes across all prior learned tasks to arrive at a final posterior on the output.

When the learner is exposed to an additional new task, a single task hypothesis is generated, along with an updated set of cross-task posterior distributions for tasks learned previously. We were able to show a number of applied examples of progressive learning using this model by the use of lifelong forests and networks. Essentially, a modified decision forest or deep network approach that makes use of such ensemble representations. We have shown that this approach is able to successfully partition rather complex decision spaces efficiently (Figure 4), outperforming other state of the art methods.

**Figure 4:**
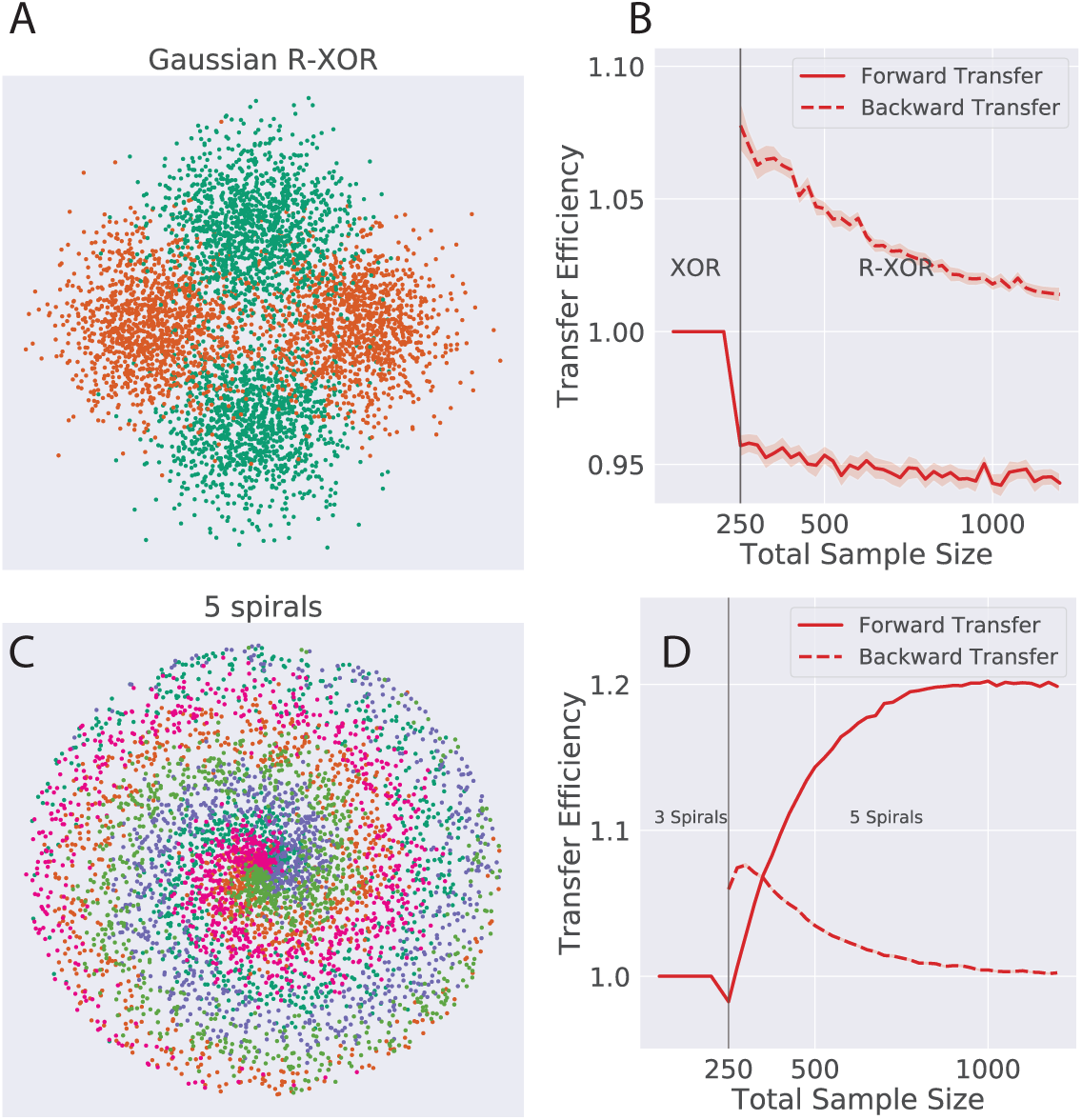
Examples of the forward and backward transfer of information in learned decision tasks. **A.** Graphical distribution of R-XOR, an XOR (’exclusive or’ logical function) space that has orthogonal decision boundaries to XOR. B. Lifelong forests showed positive forward transfer efficiency and ’graceful’ forgetting. **B**. A spiral task distribution with five classes. We created a similar spiral distribution with three classes. The five class problem has a finer optimal partition. **D**. Partition (decision separation) of the two classes using lifelong forests. Reproduced from [31].

For any neurobiological computational element that might implement progressive learning, such as a neural circuit or even an individual neuron (especially given how computationally complex and sophisticated an individual neuron can be, see for example [4, 11]), the transfer of information between each of the composite parts of the learning task hypothesis will be bounded by physiological constraints. For example, changes in firing rates or signaling routes due to learning would be bounded by a variant of the optimized refraction ratio theorem, which proves the theoretically derived ideal for efficient signaling in a geometric network in the context of the refraction ratio. Application of the theorem sets neurobiologically realistic limits on the rate of progressive transfer learning. This notion is not part of the original model. But one can construct an extension of the model that accommodates the fundamental structure-function constraint. For example, possibly through sequential tasks, where the ensemble of representations corresponds to snapshots of a system over a time windows, such that each window of time provides constraints about the extent to which the representation could change. Furthermore, we can learn new things by theoretically and numerically (computationally) analyzing the extended construction that follows the neurobiologically imposed constraints that we would not be able to otherwise.

Two final comments. First, because the competitive-refractory dynamics model is derived from direct considerations of the structure-function constraint and the anatomical and neurobiological considerations responsible for it, a detailed analyses of these ideas will likely facilitate any attempt at experimental validation of progressive learning in a neurobiological system. For example, can distinct populations of neurons be identified as voters or deciders in response to specific tasks? It has been proposed in at least one model that interacting populations of neurons organized into coupled ’modules’ can evaluate competing sensory information [22]. The dispersion between modules encodes a confidence measure that the brain can take advantage of in using that information to make subsequent decisions. Other recent experimental work suggests that both single-neuron encoding and distributed encoding schemes are used by the brain to vote on and make decisions about a task [7]. When the brain is initially processing a new input or piece of data, many neurons are polled in order to generate a good prediction of what the decision should be. But the closer the brain gets to actually making a decision, the neurons converge and begin to agree on what the right decision should be. Eventually, as a consensus is reached, each neuron on its own is maximally predictive. In other words, while the voting and deciding population of neurons is initially heterogeneous, convergence results in a collective consensus. At that point, polling almost any individual neuron would tell you the decision the brain has come to. It is a form of neurobiological partitioning.

Second, other properties associated with the mathematical ideas behind progressive learning may provide additional insights about the neurobiological mechanisms that implement it. For example, one of the canonical theorems about the composition of functions states that to preserve distinctiveness from an input to an outputs when there is an intermediate map (i.e. representation) between the input and output, there must be distinctiveness preserved in the mapping from the input to the intermediate representation, but need not be in the mapping from that intermediate representation to the output. In the context of progressive learning, this implies that as long as distinctiveness is preserved from the inputs to the their internal representations in the neural system through the transformer, distinctiveness will also be preserved in the mapping from posterior distributions to the outputs by the deciders. But need not be distinct in the mapping by voters from internal representations to posteriors. This is insightful because it means that there may exist neurobiological processes or set of algorithms that map different inputs to the same internal representations, thus conserving computational and cellular resources, something that could be experimentally testable.

## 4 Towards a systems understanding of the brain

### 4.1 Intersection with machine learning and artificial intelligence

Beyond neuroscience itself, achieving a systems engineering and theoretical understanding of the brain offer a path forward for pushing the state of the art towards the next generation of machine learning and artificial intelligence. Given sufficient convergence between experimental models (e.g. brain organoids) and data on the one hand, and theoretical and computational ideas on the other, there is an opportunity to expand the impact of neuroscience on machine learning beyond its current contributions. A systems neuroscience approach offers an opportunity to advance machine learning toward functionality, adaptability, and properties closer to animals and humans. Continued progress in other mathematical and statistical applications, as well as hardware, that have nothing to do with the brain will also drive the methods forward. But nature has a significant head start on us, and the biological brain certainly has a lot to teach us and contribute.

Turning the tables around, theoretical machine learning has the potential to inform the discovery of brain algorithms. For example, the study of convolution neural networks (CNN’s) and graph neural networks (GNN’s). However, we propose that such an approach will be successful if the structural and mathematical design and set up of the artificial networks being investigated are also appropriately constrained by the physical conditions imposed on the neurobiology.

### 4.2 Other models and data: The advantages of an inclusive approach

Throughout this paper we have attempted to build a case for the requirement that any computational model or theory the makes claims about its relevance to how the brain works must be able to account for the fundamental structure-function constraint. And because of their putative relevance to human brain structure and function, we also argued that human-derived brain organoids could be an experimental model of the brain capable of guiding the development and testing of constraint-based mathematical models. One of the unique features of organoids is their ability to bridge mechanistic neurobiology with higher level cognitive properties and clinical considerations in the intact human. The kind of parallel basic neurobiology-clinical trial paradigm we are pursing in the autism study, for example, offers a potential paradigm shift in how mechanistic neurobiology is viewed in context with clinical effects and outcomes.

While direct relevance to the human brain will be variable, other experimental models offer the opportunity to ask specific questions that take advantage of each model’s unique properties which organoids might not share. There are likely aspects of neurobiology that invariant to taxa, such that considering them in the context of what also happens in an organoid could establish an abstraction about the wetware. For example, the deterministic and reproducible connectivity of C. elegans across individuals, or the opportunity to observe and measure the physiology and metabolic properties of intact zebra fish during behavioral experiments. Rigorous constraint-based mathematical models that integrate data across experimental models offer the opportunity to generate insights about how the brain works that no single approach by itself can produce. It is the epitomy of a collective effort aimed at achieving a systems engineering understanding of the brain. It offers an opportunity to view neurological, neurodevelopmental, and psychiatric disorders in a new holistic, and even personalized, way that might produce clinical options that do not yet exist.

## Appendix

### The Hodgkin Huxley model

Consider the Hodgkin Huxley model of a neuron. Current flows in the cable equation are computed along the dominant one dimensional *x* direction of the neuron as a function of time. The membrane biophysics, due to the effects of the participating sodium and potassium ion channels, completely determines the passive spatial and temporal decay of potentials along the membrane. Explicitly, the change in membrane potential as a function of *x* given appropriate boundary conditions is solved to be

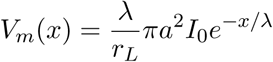

for an initial current step *I*_0_ at *x* = 0, membrane radius a, and parametrized membrane resistance *r*_*L*_. *λ* represents the space constant, which is the distance over which the potential has decayed to 1/*e* its original amplitude. This is a critical consideration for the spatial and temporal summation of post synaptic potentials along the dendritic tree towards reaching the threshold potential necessary to trigger an action potential at the axon hillock. This spatial-temporal dependency is equally explicit in the Hodgkin Huxley equations. The membrane potential dynamically depends on the spatial variable *x* and propagation latencies at times *t*. Assuming standard notations and Ohmic ionic currents, as is typical, the equation that balances the various components of the membrane potential is given by:

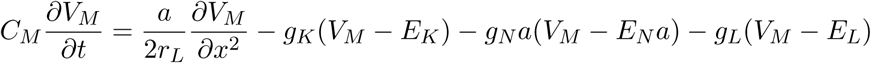

In the solution to its equations, the gating variables ultimately determine the temporal kinetics of the action potential. If the membrane refractory state associated with the recovery of sodium channel inactivation is taken into consideration, the conduction velocity of the traveling action potential can be derived. This reflects the rate at which information can be communicated or transferred between neurons. We again refer the reader to the standard treatments of the Hodgkin Huxley model for full details [1, 5, 9, 16].

### Competitive-refractory dynamics model

Assume a directed graph *G* = *(V, E*) such that the edge *e*_*ij*_ is the directed edge from vertex *v*_*i*_ to vertex *v*_*j*_. Define the subgraph *H*_*j*_ as the (inverted) tree graph that consists of all vertices *v*_*i*_ with directed edges into *v*_*j*_. Across an entire network with |*V*| vertices, there will be the same number of *H*_*j*_ subgraphs (i.e. one for each node in the network). For each *H*_*j*_, the model fractionally summates discrete signaling events traveling at a finite speed *s*_*ij*_ on edges *e*_*ij*_ until a threshold is reached and *v*_*j*_ fires. But because the model explicitly considers geometric networks, the offsets in the latencies of arriving signaling events at *v*_*j*_ have a profound effect on the fractional summation. In addition, weights associated with each arriving signal progressively decay over time, and therefore progressively less contribute less to the summation at each subsequent time step. This adds a further important dynamical variable to the fractional summation.

More formally, at an observation time *T*_*o*_ a signal from any *v*_*i*_ may be traveling part way along an *e*_*ij*_ with length |*e*_*ij*_| at a speed *s*_*ij*_, effectively shortening the absolute signal latency *τ*_*ij*_· along *e*_*ij*_. Or it may be delayed in signaling if *v*_*i*_ signals some time after *T*_*o*_, effectively lengthening *τ*_*ij*_. Each vertex *ν*_*i*_ ∈ *H*_*j*_(*ν*_*i*_) is independent of every other vertex in the subgraph *H*_*j*_ (*v*_*i*_). And as such different temporal offsets are to be expected depending on how far along a discrete signaling event is in its propagation along their respective edge *e*_*ij*_. Thus, the distance that a discrete signal initiated by each *v*_*i*_ has traveled along its edge at *T*_*o*_ is taken into account by defining a temporal offset for *τ*_*ij*_, an effective shortening or lengthening of *τ*_*ij*_ relative to *T*_*o*_ as

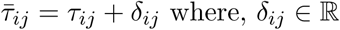

The second key concept is the consideration of an absolute refractory period for the vertex *ν*_*j*_ by *R*_*j*_. This reflects the internal dynamics of *ν*_*j*_ once a signaling event activates it. For example, the amount of time the internal dynamics of *ν*_*j*_ requires to make a decision about an output in response to being activated, or some reset period during which it cannot respond to subsequent arriving input signals. There are no restrictions on the internal processes that contribute toward *R*_*j*_. Since nothing is instantaneous, the assumption is that once a node activates, it becomes refractory for a period *R*_*j*_ that progressively decreases towards zero (i.e. a state of not being refractory anymore). Similar to the consideration of 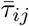, *T*_*o*_, there is likely to be a temporal offset between when signals from each *ν*_*i*_ arrive and how far along *ν*_*j*_ is in its recovery from its refractory period. Furthermore, each *v*_*i*_ ∈ *H*_*j*_(*v*_*i*_) is statistically independent from every other *ν*_*i*_, so that the amount of temporal offset for each *ν*_*i*_ vertex signaling *ν*_*j*_ will be different. Mathematically the model accounts for this in the following way. Let *ϕ*_*j*_ represent a temporal offset from *R*_*j*_, such that at *T*_*o*_

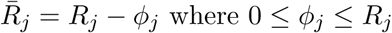

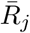 is the effective refractory period. It reflects the amount of time remaining in the recovery from *R*_*j*_ at the observation time *T*_*o*_.

The refradtion ratio, the ratio between the relative refractory period and effective latency (relative to an observation time *T*_*o*_), is then written as

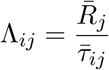

This expression is key, because it describes a local rule (at the individual node scale, i.,e. *H*_*j*_ for every vertex *ν*_*j*_) that governs the global dynamics of the network. Although beyond the scope of this paper, by creating a well ordered set of refraction ratios one can compute the causal order in which incoming signaling events trigger the activation of any node *ν*_*j*_ for all participating *ν*_*i*_ nodes in *H*_*j*_.

The last important consideration for our purposes here, is that a theoretical analysis results in the derivation of a set of (upper and lower) bounds that constrained a mathematical definition efficient signaling for a structural geometric network running the dynamic model. This produced a proof of the optimized refraction ratio theorem. If we let *G* = *(V, E*) represent a geometric network consisting of subgraphs *H*_*j*_ (*ν*_*i*_). For each *ν*_*i*_*ν*_*j*_ vertex pair with a signaling speed *s*_*ij*_ between *ν*_*i*_ and *ν*_*j*_, the optimal refraction ratio [Λ_*ij*_]_*opt*_ at an observation time *T*_*o*_ is bounded by

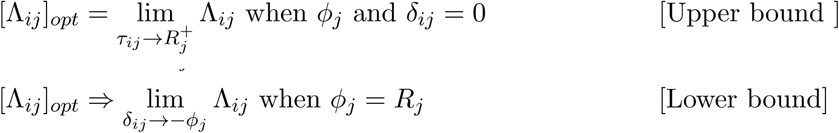

Given these bounds then, the optimized refraction ratio is

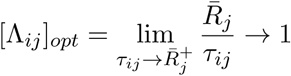

We refer the reader to the original paper for full details and proofs [26].

### Progressive transfer learning model

Progressive learning is an approach that supports transfer learning across tasks in both a forward and backward direction [31]. The core of their argument relies on being able to evaluate a hypothesis 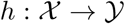. For the single task case, *h* is decomposed into three parts, each with a successive purpose: *h*(⋅) = *w* o *v* o *u*(⋅). Given a *χ*-valued input the transformer u maps the input into an internal representation 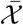. Following the construction of an internal representation to the input, the voter 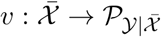 maps the transformed input data into a posterior distribution on a response space 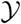. In the last step, the decider *w* maps 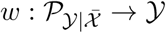 to an output 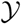. The model’s core insight and innovation is the construction of ensemble representations, or grouping, of learned transformations across tasks, which can then be used by deciders on new tasks. In other words, the transfer of learned information between different tasks contributes to learning new tasks. To achieve this, assume *n* input samples produce a set of 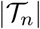 tasks in the environment 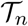. This creates 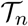 cross-task posteriors. The decider *w*_*t*_ for task t combines the votes across all prior learned tasks to arrive at a final posterior on the output 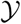. If the function that computes the combination of prior learned tasks is an average, then the hypothesis for task *t*’ *h*_*t*_ (expanding the authors’ original notation a bit) would be

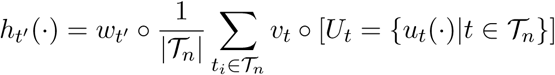

When the learner is exposed to an additional new task *s*, a single task hypothesis *h*_*s*_ = *w* o *v* o *u* to *s* is generated, along with an updated set of cross-task posteriors {*ν*_*t*_ o *u*_*s*_}_*t*∈*T*_*n*__. Note how in this notation, *t* ∈ *T*_*n*_ represents the aggregation of all prior learned tasks, i.e. *t*’ ∈ *T*_*n*_ in the above ensemble hypothesis equation *h*_*t′*_. In their original paper the authors illustrate a number of applied examples of progressive learning using this model by the use of lifelong forests and networks. Essentially, a modified decision forest approach that makes use of such ensemble representations.

For any neurobiological computational element that implements this model, such as a neural circuit or even an individual neuron, the transfer of information between each of the composite parts of hypothesis *h* will be bounded by a variant of the optimized refraction ratio theorem. This sets physiologically realistic limits on the rate of progressive transfer learning that is not part of the original model itself. The conjecture is that there exits (a provable) extension of the model to accommodate the fundamental structure-function constraint.

Assume an ensemble *ξ*_*υ*_ of neural elements that is responsible for mapping an input *u*_*t*_ or environment 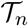 to an encoding representation *U*_*t*_ -which could be an external sensory input or an internal representation of some element of an inherent internal computation. Leveraging the theoretical framework of the competitive-refractory dynamics model, let the relative time associated with the internal transfer of information (message passing) within the parts of *ξ*_*υ*_ and computational time at an observation time *T*_*o*_ be 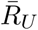. Let 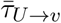 be the relative time it takes for information transfer from transformers to voters as a function of the anatomical and physiological processes that invoke it, also at *T*_*o*_. We modify our original notation such that 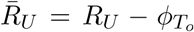. *R*_*u*_ reflects the the dynamical requirements associated with a specific encoding scheme of the input. 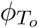 modulates the remaining time associated with *Ru* given when one begins to observe or measure the process, i.e. at time *T*_*o*_ relative to the history of the system. Similarly, 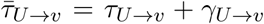, such that *γ*_*U*→*ν*_ modulates the absolute transfer time due to the physiology by what remains, referenced to the observation time *T*_*o*_ at which the system is measured. A similar set of equations exists between voters and deciders, namely,

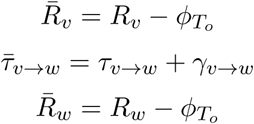

If we assume that the introduction and learning of a new task s, including its integration into the existing environment of previously learned priors, i.e. into 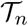, cannot occur before previously learned tasks have been similarly integrated, a reasonable requirement given physiological and causality considerations, then the optimized refraction ratio theorem set bounds on how quickly progressive learning can take place. Specifically:

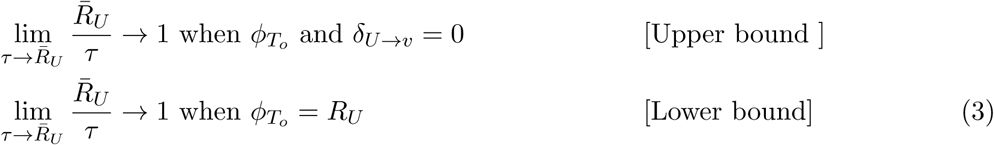

where for simplicity in notation *τ* here implies *τ*_*U*→*ν*_. Similar bounds can be derived for the mapping from voters to deciders.

Other properties associated with the mathematical ideas behind progressive learning may provide additional insights about the neurobiological mechanisms that implement it. For example, one of the canonical theorems about the composition of functions states that *If f*: *A* → *B and g*: B → *C are mappings such that g* o *f: A* → *C is injective, then f is injective.* The theorem proves that *g o f* is injective if *f* is injective, even if *g* is not. This means that to preserve distinctiveness from the mapping *A → C* there must be distinctiveness preserved in the mapping from A → B, but need not be in the mapping from *B → C*. In the context of progressive learning, this implies that as along as distinctiveness is preserved from the inputs to the their internal representations in the neural system through the transformer *u*, distinctiveness will also be preserved in the mapping from posterior distributions 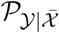 to outputs 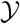 by the deciders *w*. But need not be distinct in the mapping by voters from internal representations to posteriors. This is insightful because it means that there may exist neurophysiological processes or set of algorithms that map different inputs to the same 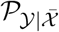, thus conserving computational and cellular resources. This is something that could be experimentally testable.

## Notes

### Competing Interest Statement

The authors have declared no competing interest.

### Summary of Updates

Fixed a number of missing references. The text was not changed from the original version.

